# Low-Density Lipoprotein Modulates Plasma Fibrin Network Architecture and Impairs Fibrinolysis

**DOI:** 10.64898/2026.08.31.748310

**Authors:** Arezoo Nameny, Austin DeSmet, Can Cai, Stephen R. Baker, Keith Bonin, Nathan E. Hudson, Brittany E. Bannish, Martin Guthold

## Abstract

Low-density lipoprotein (LDL) is a major atherogenic lipoprotein, yet its potential to directly modify the fibrin scaffold of blood clots is incompletely understood. Here, we investigated how LDL alters plasma fibrin network architecture and internal fibrinolysis across defined fibrinogen/thrombin conditions. Pooled normal human plasma was supplemented with LDL and clotted with controlled concentrations of fibrinogen and thrombin. Fibrin architecture was visualized by confocal microscopy and quantified by pore-size analysis; clot formation and lysis were monitored turbidimetrically in the presence of tissue plasminogen activator (tPA). Increasing LDL produced a pronounced reduction in fibrin-network pore size across the tested fibrinogen/thrombin conditions. The LDL dependence of pore diameter was well described by a power-law relationship, D_pore = (6.54 ± 0.11)[*LDL*]^−0.12 ± 0.02^ , (R^2^ = 0.90), with a significant negative LDL exponent (p = 4 × 10^−5^). Increasing LDL also prolonged clot lysis time and altered turbidity kinetics. These findings extend epidemiologic and clinical associations between ApoB-containing lipoproteins and hypofibrinolytic clot phenotypes by demonstrating, in a controlled plasma system, that LDL itself can modify fibrin network architecture and fibrinolytic susceptibility. The results support a structure-function role for LDL within the fibrin biomaterial and motivate direct tests of LDL incorporation, protofibril packing, fibrinolytic-protein binding, and single-fiber mechanics.

## 1. Introduction

Fibrin is the load-bearing protein scaffold of a blood clot and forms a hierarchical biomaterial whose function depends on molecular assembly, individual-fiber properties, and network architecture. Thrombin-mediated conversion of fibrinogen to fibrin produces branched fiber networks whose pore dimensions and fiber morphology vary with the biochemical conditions of polymerization. Recent quantitative work has shown nonlinear dependence of plasma and purified fibrin architecture on fibrinogen and thrombin concentrations, emphasizing that the biochemical environment during polymerization establishes the structural baseline on which other plasma constituents act [1-2].

Fibrin architecture is relevant to fibrinolysis, but structural changes alone do not necessarily determine the rate of internal clot lysis. In our recent controlled clot study, increasing both fibrinogen and thrombin produced marked changes in fibrin network architecture, whereas internal clot lysis time was governed primarily by fibrinogen concentration and was largely independent of thrombin-mediated structural variations and clot-formation kinetics [2]. These findings demonstrate that changes in network architecture do not necessarily translate into corresponding changes in internal fibrinolytic susceptibility. At the single-fiber scale, fibrin fibers possess distinctive mechanical properties, including high extensibility and strain-dependent stiffness [3], and mechanical strain can markedly reduce the susceptibility of individual fibers to lysis [4]. Fiber diameter and prestrain have also been linked to distinct single-fiber lysis behaviors [5], while recent work indicates that protofibril packing within individual fibers can further modulate fibrinolytic behavior [6]. At the network scale, pore dimensions and network density may influence the transport and redistribution of fibrinolytic proteins, and fibrinolysis itself dynamically remodels network pores [7]. However, because the structural variations in that study were generated by varying fibrinogen concentration at a fixed thrombin concentration, the independent contribution of thrombin-mediated changes in network architecture to internal fibrinolysis was not assessed. In our recent study, systematic variation of both fibrinogen and thrombin demonstrated that thrombin-induced changes in network architecture did not result in corresponding changes in internal clot lysis time [2]. Thus, the biochemical composition and material properties of the clot, rather than network architecture alone, may be critical determinants of internal lysis. This distinction provides an important framework for testing whether LDL alters fibrinolysis solely through structural remodeling or through additional mechanisms.

Low-density lipoprotein (LDL) is an ApoB-100-containing particle that transports cholesterol in the circulation and plays a central role in atherosclerosis. Native LDL is a nanoscale lipid-protein assembly consisting of a hydrophobic lipid core surrounded by a surface monolayer containing phospholipids, unesterified cholesterol, and a single apoB-100 molecule [8]. Recent cryo-electron microscopy and integrative structural analysis of human LDL have provided a detailed molecular model of apoB100 organization on the LDL particle, emphasizing the extensive amphipathic protein-lipid interface presented by LDL [9]. The established cardiovascular effects of LDL are generally discussed in the context of arterial-wall retention and atherogenesis; however, accumulating evidence links ApoB-containing lipoproteins to fibrin clot phenotype. Proteomic analyses demonstrate that plasma clots retain numerous nonfibrin proteins, including apolipoproteins [10]. Clinical and population studies have associated higher LDL-C and ApoB levels with prothrombotic fibrin clot characteristics, including denser or less permeable clot networks and prolonged clot lysis [11–13]. Furthermore, intensive LDL-C lowering has been associated with increased clot permeability and shorter clot lysis time in patients with coronary artery disease [12]. Broader reviews likewise support reciprocal links among lipids, fibrin clot phenotype, and thrombotic biology [14–15].

Related lipoprotein studies further support the possibility that lipid-protein particles can modify fibrin function, although LDL must be distinguished mechanistically from lipoprotein(a) [Lp(a)]. Lp(a) is an ApoB-containing particle with an additional apolipoprotein(a) component and has recently been shown to alter ex vivo clot formation, fibrin architecture, and clot lysis [16]. Likewise, lipid-associated plasma constituents can modify fibrin as a biomaterial: fatty acid-albumin conjugates were recently reported to alter fibrin network structure, mechanics, and degradation [17]. Together, these observations raise a distinct mechanistic question: can LDL itself directly perturb fibrin assembly and fibrinolytic susceptibility, independent of the clinical covariates that accompany dyslipidemia?

The present study addresses this question using a controlled plasma-clot system in which LDL concentration was varied across defined fibrinogen/thrombin conditions. We used confocal microscopy to quantify fibrin-network pore size and turbidimetry to characterize clot formation and internal lysis. We further tested whether a power-law relationship could describe network pore size as a function of LDL concentration. Building on our previous finding that fibrinogen, rather than thrombin-mediated structural variation, was the primary determinant of internal clot lysis time [2], we specifically asked whether LDL-dependent structural remodeling is accompanied by a change in fibrinolytic susceptibility. We hypothesized that increasing LDL would alter fibrin network architecture and prolong internal clot lysis. Given our previous finding that network structural changes do not necessarily predict internal clot lysis time, we further sought to determine whether LDL-induced changes in structure and fibrinolysis were associated, without assuming a direct causal relationship between them [2].

## 2. Materials & Methods

### 2.1 Material and plasma preparation

Pooled normal human plasma was obtained from George King Bio-Medical, Inc. (Product No. 0020, Lot 7716, Overland Park, KS; stock concentration, 3.25 mg/mL). Plasma was aliquoted and stored at −80 °C. Before use, aliquots were thawed at 37 °C for 10 min. Baseline LDL-C in the pooled plasma was measured using an LDL cholesterol assay kit (Crystal Chem, Catalog No. 80069) and was reported as 0.7 mg/mL. Human LDL isolated from plasma was obtained from Medix Biochemica USA, Inc. (Catalog No. 360-10, Lot 02K9502), aliquoted, stored at −80 °C, and warmed at 37 °C for 10 min before use. Human thrombin was obtained from Enzyme Research Laboratories, South Bend, IN (Catalog No. HT 1002a, Lot HT 6252A). Alexa Fluor 488-conjugated fibrinogen (Thermo Fisher Scientific, Catalog No. F 13191, Lot 2604402) was aliquoted and stored at −80 °C. Before use, fluorescent fibrinogen was warmed to 37 °C and briefly centrifuged (10 sec.) to remove visible aggregates. Tris-buffered saline (TBS) contained 20 mM Tris and 150 mM NaCl, pH 7.4.

### 2.2 Fluorescent plasma-clot preparation

Plasma was diluted in TBS to generate target fibrinogen concentrations of 0.73, 1.45, and 2.5 mg/mL. Alexa Fluor 488-conjugated fibrinogen was added at 1.5–10.5% of the total fibrinogen concentration, with the labeling fraction adjusted according to fibrinogen concentration. These labeling fractions were selected based on preliminary optimization experiments to provide sufficient fiber fluorescence for quantitative confocal analysis while minimizing fluorescent aggregates and labeling-associated imaging artifacts observed at higher labeling fractions [2]. Exogenous LDL-C was added at 0.5, 1.0, 2.0, and 3.0 mg/mL. Clotting was initiated with thrombin and CaCl2 to yield final thrombin concentrations of 0.1, 0.2, or 1.0 U/mL and a final CaCl2 concentration of 20 mM. The fibrinogen/thrombin pairs used for the LDL matrix were 0.73 mg/mL/0.1 U/mL, 1.45 mg/mL/0.2 U/mL, and 2.5 mg/mL/1.0 U/mL. Immediately after mixing, 30 μL was transferred to channel slides (ibidi μ-Slide VI 0.4), protected from light, and allowed to polymerize for 2 h at room temperature.

### 2.3 Confocal microscopy and network analysis

Fibrin networks were imaged using a Zeiss LSM 710 laser-scanning confocal microscope with a 40x oil-immersion objective and 488-nm excitation. For each condition, three independent channels were prepared, and three locations within each channel were imaged, yielding a total of nine image stacks per condition. Image stacks covered an x-y area of 212 μm × 212 μm with a z-stack depth of 15 μm, acquired in 33 slices at 0.47 μm intervals. Images were collected at 8-bit resolution. The master gain was adjusted to visualize fibers while minimizing background noise. Maximum-intensity projections were generated for visualization. Network pore size was quantified using the same image-analysis procedure as in our previous fibrin-network study [2].

### 2.4 Bubble analysis

Fibrin-network pore dimensions were quantified from individual confocal z-slices using a custom MATLAB-based image-analysis algorithm previously described [18]. Images were exported as multi-page TIFF files and binarized using an intensity threshold defined as three times the mean intensity of each slice. The algorithm identified fiber-free regions and iteratively fitted maximally sized circles, “bubbles”, within these regions. Bubbles intersecting fibrin fibers or image boundaries, as well as those overlapping larger accepted bubbles, were excluded. The diameters of the retained bubbles were used as a quantitative measure of fibrin-network pore size, with larger diameters corresponding to larger fiber-free regions.

### 2.5 Turbidimetric clot formation and internal fibrinolysis

Clot formation and internal fibrinolysis were monitored turbidimetrically using a SpectraMax microplate reader by measuring absorbance at 405 nm every 7 s at 37 °C for 3 h. Reactions were performed in 96-well microplates (Corning flat clear-bottom white polystyrene, tissue-culture treated) with a final reaction volume of 100 μL per well. Three fibrinogen/thrombin concentration combinations were examined: 0.73 mg/mL fibrinogen with 0.1 U/mL thrombin, 1.45 mg/mL fibrinogen with 0.2 U/mL thrombin, and 2.5 mg/mL fibrinogen with 1.0 U/mL thrombin. Human tissue-type plasminogen activator (tPA; Sigma-Aldrich) was added to the fibrinogen solution before initiation of clotting to induce internal fibrinolysis. Based on our previous findings that fibrinogen concentration is a primary determinant of internal clot lysis time [2], the tPA concentration was optimized for each condition to allow both clot formation and complete fibrinolysis to be captured within the 3-h measurement period. Final tPA concentrations were 26, 45, and 110 ng/mL for the 0.73/0.1, 1.45/0.2, and 2.5/1.0 mg/mL fibrinogen, U/mL thrombin conditions, respectively. Within each condition, tPA concentration was held constant across all LDL concentrations; therefore, LDL-dependent changes in clot formation and lysis were evaluated within each fibrinogen/thrombin/tPA condition rather than by direct comparison among the three condition sets. Clot lysis time (CLT) was defined as the time interval between the point at which absorbance reached 50% of the maximum during clot formation and the point at which it decreased to 50% of the maximum during clot lysis [19].

### 2.6 Statistical analysis

All statistical analyses were performed in R version 4.5.1 using the base stats package. The dependence of fibrin-network pore size on fibrinogen, thrombin, and LDL concentrations was initially evaluated using a multiplicative power-law model, D_pore = *k*([*Fgn*]^*α*^ × [*Thr*]^*β*^ × [*LDL*]^γ^) where D_pore represents pore diameter, *k* is the scaling coefficient, and *α, β*, and γ represent the fitted exponents for fibrinogen, thrombin, and LDL concentrations, respectively. Model parameters were estimated by nonlinear least-squares regression by minimizing the sum of squared residuals between the observed and model-predicted pore diameters. Standard errors were obtained from the covariance matrix of the fitted model, and two-sided t-tests were used to assess whether individual parameter estimates differed significantly from zero.

In the full three-variable power-law model, the fitted exponents for fibrinogen and thrombin were not statistically distinguishable from zero after accounting for LDL concentration, whereas the LDL exponent remained significant. This indicates that, within the concentration ranges and experimental conditions examined here, LDL concentration accounted for the dominant concentration-dependent variation in the measured response. A reduced power-law model containing LDL as the sole concentration-dependent predictor was therefore fit subsequently. D_pore = *k*[*LDL*]^γ^.

For fibrinolysis, clot lysis time (CLT) was analyzed separately within each fibrinogen/thrombin/tPA condition as a function of total LDL concentration using a power-law model, *CLT* = *k*[*LDL*]^δ^, where *k* is the scaling coefficient and δ is the fitted exponent describing the dependence of CLT on LDL concentration. Separate nonlinear least-squares fits were performed for each of the three fibrinogen/thrombin/tPA conditions. Parameter standard errors were obtained from the covariance matrix of each fitted model, and two-sided t-tests were used to determine whether the LDL exponent differed significantly from zero. Goodness of fit was assessed using R^2^. Fibrinogen and LDL concentrations were expressed in mg/mL and thrombin concentration in U/mL. LDL concentration was reported on an LDL-C basis.

## 3. Results

### 3.1 LDL reduces fibrin-network pore size across fibrinogen/thrombin conditions

Representative maximum-intensity projections revealed progressive remodeling of plasma fibrin networks with increasing LDL concentration (Fig. 1). Across all three fibrinogen/thrombin condition sets, increasing LDL was associated with visually smaller inter-fiber spaces. Quantitative pore-size analysis, based on three independently prepared samples with three confocal stacks acquired per sample (nine stacks per condition), confirmed a concentration-dependent decrease in pore diameter with increasing LDL concentration (Fig. 2).

### 3.2 Pore diameter follows a power-law dependence on LDL concentration

Quantitative analysis of the confocal images demonstrated a progressive decrease in fibrin-network pore diameter with increasing LDL concentration (Fig. 2). To assess the independent contributions of fibrinogen, thrombin, and LDL concentration, pore diameter was initially fit to the three-variable power-law model D_pore = *k*([*Fgn*]^*α*^ × [*Thr*]^*β*^ × [*LDL*]^γ^). The fitted exponents were *α* = − 0.05 ± 0.09 for fibrinogen (p = 0.55), *β* = − 0.02 ± 0.05 for thrombin (p = 0.65), and γ = − 0.1 ± 0.01 for LDL (p = 2.3 × 10−5). The full model explained 87% of the observed variation in pore diameter (R^2^ = 0.87). The fibrinogen and thrombin exponents were not statistically distinguishable from zero, whereas the LDL exponent was significant. A reduced power-law model containing LDL as the sole concentration-dependent predictor was therefore subsequently fit to the data. Pore diameter decreased systematically with increasing LDL concentration and was well described by the power-law relationship D_pore = (6.54 ± 0.11)[*LDL*]^−0.12 ±^ 0^.02^, The model explained 90% of the observed variation in pore diameter (R^2^ = 0.90). The negative LDL exponent was significantly different from zero (p = 4 × 10−5), demonstrating a significant inverse relationship between LDL concentration and fibrin-network pore diameter over the concentration range investigated. LDL concentrations used for quantitative analysis were expressed on an LDL-C basis, as described in Materials and Methods.

**Figure 1.**
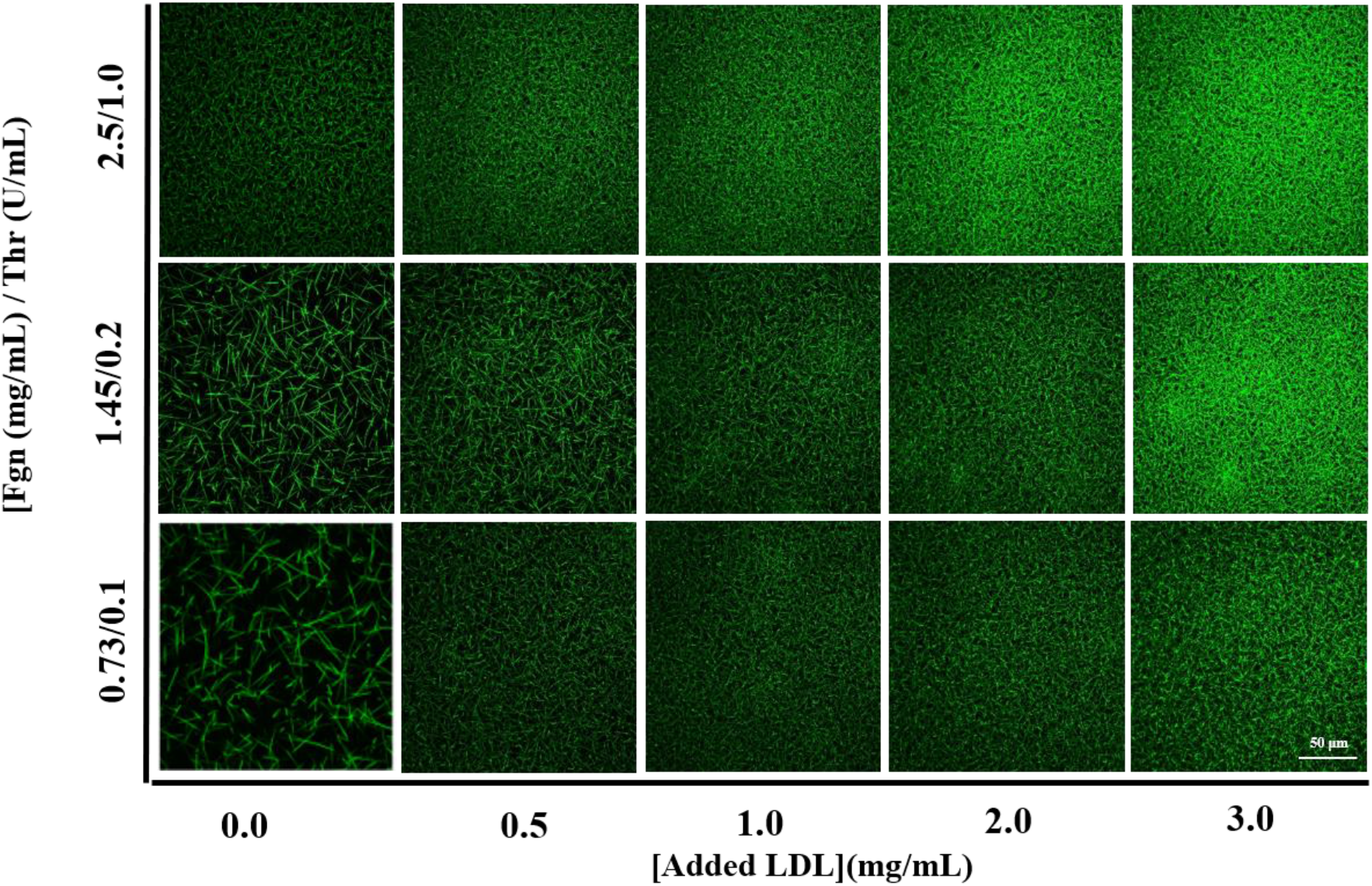
LDL-dependent remodeling of plasma fibrin network architecture. Representative maximum-intensity projections of confocal z-stacks showing plasma fibrin networks formed at varying LDL concentrations and fibrinogen/thrombin conditions. Images are arranged as a 3 × 5 matrix, with increasing added LDL concentration across columns and fibrinogen/thrombin conditions across rows. The fibrinogen/thrombin conditions were 0.73 mg/mL/0.1 U/mL, 1.45 mg/mL/0.2 U/mL, and 2.5 mg/mL/1.0 U/mL, respectively. Added LDL concentrations are expressed on an LDL-C basis and represent 0.0, 0.5, 1.0, 2.0, 3.0 mg/mL. Increasing LDL was associated with progressively smaller inter-fiber spaces across the three fibrinogen/thrombin conditions. Scale bar, 50 μm.

**Figure 2.**
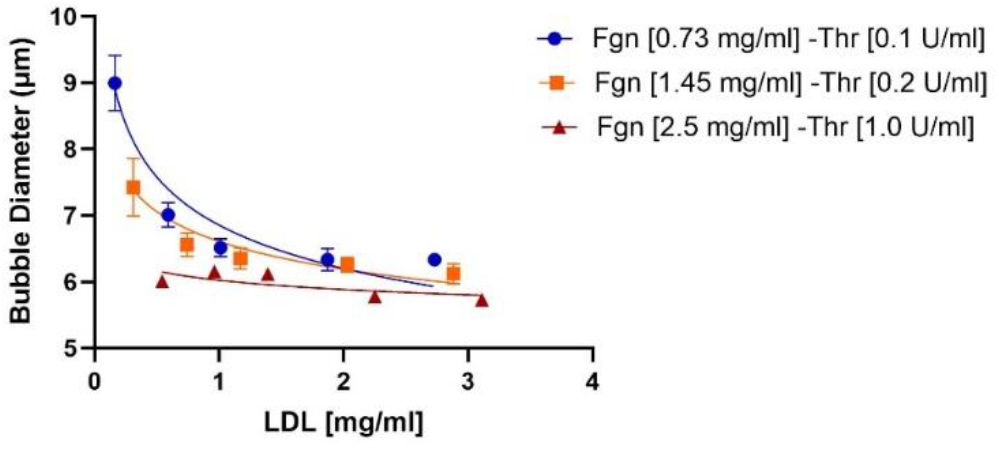
LDL-dependent decrease in plasma fibrin-network pore diameter. Quantitative pore-diameter analysis of plasma fibrin networks as a function of LDL concentration. Pore diameter was determined from confocal images using the bubble-analysis method. The solid line represents the nonlinear least-squares fit to the reduced power-law model *D* = *k*[*LDL*]^γ^. The fitted relationship was D_pore = (6.54 ± 0.11)[*LDL*]^−0.12 ±^ 0^.02^, with R^2^ = 0.90. The LDL exponent was significantly different from zero (p = 4 × 10−5). LDL concentrations are expressed on an LDL-C basis and represent total LDL-C (baseline plasma LDL-C plus supplemented LDL-C). [n = 9 per condition].

### 3.3 Increasing LDL prolongs internal clot lysis time

LDL produced a concentration-dependent prolongation of internal clot lysis across all three fibrinogen/thrombin/tPA conditions (Fig. 3). Increasing LDL concentration was associated with delayed clot dissolution in the turbidity profiles (Fig. 3A–C). The fibrinogen-free controls remained stable over time, although baseline absorbance increased with increasing LDL concentration, indicating an LDL-dependent contribution to turbidity at 405 nm in the absence of fibrin formation (Fig. 3D). Quantitative analysis demonstrated increasing clot lysis time (CLT) with increasing LDL concentration (Fig. 3E). The LDL dependence of CLT was well described by the power-law relationship *CLT* = *k*[*LDL*]^δ^ within each experimental condition (Table 1). For clots containing 0.73 mg/mL fibrinogen and 0.1 U/mL thrombin, the fitted exponent was δ = 0.187 ± 0.006, (p = 3.33 × 10−^13^, R^2^ = 0.99). For 1.45 mg/mL fibrinogen and 0.2 U/mL thrombin, δ = 0.261 ± 0.008, (p = 5.58 × 10−^14^, R^2^ = 0.99), while for 2.5 mg/mL fibrinogen and 1.0 U/mL thrombin, δ = 0.253 ± 0.03, (p = 1.17 × 10−^6^, R^2^ = 0.86). Thus, the positive and statistically significant LDL exponents across all three conditions demonstrate a consistent increase in CLT with increasing LDL concentration.

**Table 1.** Power-law fits describing the dependence of clot lysis time on LDL concentration. Clot lysis time was fit separately within each fibrinogen/thrombin/tPA condition using *CLT* = *k*[*LDL*]^δ^. LDL concentration was expressed on an LDL-C basis.

| Parameters for |  | Value | Std. Error | P-Value | R<sup>2</sup> |
| --- | --- | --- | --- | --- | --- |
| 0.73 mg/mL fgn, 0.1 U/mL thr, tPA 26 ng/mL | k | 6.138e3 | 35 | <2e-16 | 0.99 |
| | $\delta$ | 0.187 | 0.006 | 3.33e-13 | |
| 1.45 mg/mL fgn, 0.2 U/mL thr, tPA 45 ng/mL | k | 5.747e3 | 35 | <2e-16 | 0.99 |
| | $\delta$ | 0.261 | 0.008 | 5.58e-14 | |
| 2.5 mg/mL fgn, 1.0 U/mL thr, tPA 110 ng/mL | k | 3.627e3 | 83 | 1.81e-15 | 0.86 |
| | $\delta$ | 0.253 | 0.03 | 1.17e-06 | |

**Figure 3.**
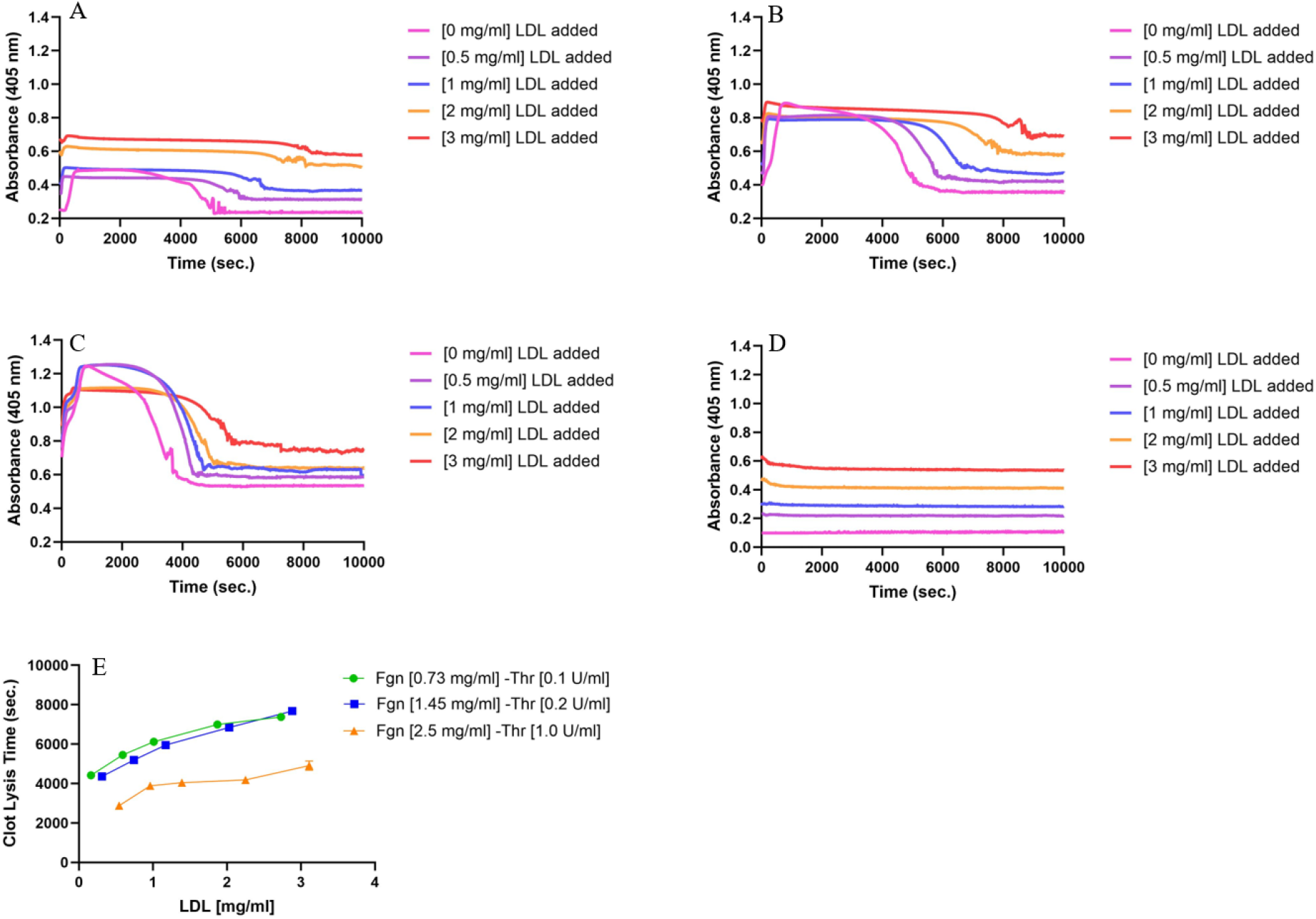
Effect of LDL concentration on plasma clot formation and internal fibrinolysis. Representative turbidity profiles showing clot formation and lysis at increasing LDL concentrations under three fibrinogen/thrombin/tPA conditions: (A) 0.73 mg/mL fibrinogen, 0.1 U/mL thrombin, and 26 ng/mL tPA; (B) 1.45 mg/mL fibrinogen, 0.2 U/mL thrombin, and 45 ng/mL tPA; and (C) 2.5 mg/mL fibrinogen, 1.0 U/mL thrombin, and 110 ng/mL tPA. (D) Fibrinogen-free control containing increasing LDL concentrations and all other reaction components, including thrombin, tPA, CaCl*2*, and buffer, monitored at 405 nm. (E) Clot lysis time (CLT) as a function of total LDL-C concentration for the three fibrinogen/thrombin/tPA conditions. LDL concentrations are expressed on an LDL-C basis and represent total LDL-C (baseline plasma LDL-C plus supplemented LDL-C). [n = 3 per condition].

Because different tPA concentrations were used for the three fibrinogen/thrombin conditions to obtain measurable clot formation and lysis within the 3-h experimental window, LDL-dependent changes in CLT were evaluated within each fibrinogen/thrombin/tPA condition rather than by direct comparison of absolute CLT values among the three conditions. Collectively, these results demonstrate that increasing LDL prolongs internal clot lysis across distinct fibrinogen/thrombin conditions.

## 4. Discussion

The principal finding of this study is that increasing LDL alters both the architecture and fibrinolytic behavior of plasma fibrin clots under controlled polymerization conditions. Increasing LDL was associated with a progressive reduction in fibrin-network pore size and prolongation of internal clot lysis across the fibrinogen/thrombin conditions examined. The dependence of pore diameter on LDL concentration was well described by a power-law relationship, and CLT likewise exhibited a significant positive power-law dependence on LDL within each fibrinogen/thrombin/tPA condition. Together, these findings provide controlled experimental evidence that LDL supplementation is sufficient to modify fibrin-network organization and reduce fibrinolytic susceptibility in an ex vivo plasma system.

These findings are consistent with human studies linking ApoB-containing lipoproteins to prothrombotic fibrin phenotypes. In a large population study, LDL-C and ApoB were positively associated with clot lysis time, while LDL-C was also associated with a turbidity-derived measure of clot density [13]. In patients with coronary artery disease, intensive LDL-C lowering was accompanied by improvement in fibrin clot phenotype [12]. Associations among apolipoproteins, Lp(a), and hypofibrinolytic clot properties have also been reported in severe aortic stenosis [11]. More broadly, Zheng and colleagues have synthesized evidence supporting links between dyslipidemia, ApoB-containing lipoproteins, and fibrinolytic regulation [20,21]. Although these clinical studies establish a biologically relevant association, they cannot readily distinguish direct effects of lipoproteins on the fibrin clot from concomitant differences in fibrinogen, inflammation, medication use, and other patient-level variables. The controlled LDL-supplementation approach used here therefore provides complementary evidence that LDL itself can contribute to changes in fibrin clot architecture and lysis.

The effects observed here should also be distinguished from those attributed to Lp(a). Although LDL and Lp(a) both contain ApoB100, Lp(a) additionally contains apolipoprotein(a), which shares structural homology with plasminogen and can interfere with fibrinolytic pathways. Recent work showed that Lp(a) prolongs ex vivo clot lysis through effects involving clot formation and fibrin architecture, while isolated apo(a) exhibited distinct antifibrinolytic activity [16]. The present findings demonstrate that LDL, an ApoB-containing particle lacking apo(a), can also alter fibrin-network architecture and prolong clot lysis. Thus, lipoprotein-associated modulation of fibrin clot properties may not be restricted to the classical apo(a)–plasminogen mechanism.

An important question is whether the reduction in pore size observed with increasing LDL is responsible for the accompanying prolongation of clot lysis. The present data demonstrate parallel LDL-dependent changes in these two outcomes but do not establish a direct causal relationship between them. This distinction is particularly important in light of our previous controlled plasma-clot study, in which increasing fibrinogen and thrombin both produced substantial changes in fibrin-network architecture, whereas internal clot lysis time was governed primarily by fibrinogen concentration and was largely independent of thrombin-mediated structural variations and clot-formation kinetics [2]. These findings demonstrated that changes in network architecture do not necessarily translate directly into corresponding changes in internal fibrinolytic susceptibility.

Nevertheless, LDL-induced network remodeling may contribute to impaired fibrinolysis through mechanisms that are not reproduced by thrombin-induced structural changes. During internal fibrinolysis, pore dimensions may influence the effective transport, redistribution, and rebinding of tPA, plasminogen, and plasmin within the fibrin network, while fibrinolysis itself dynamically remodels the network through progressive pore expansion [7]. Thus, the smaller pore dimensions observed with increasing LDL could modify the spatial environment in which fibrinolytic reactions occur. However, because our previous results demonstrate that network structure alone is insufficient to predict internal CLT [2], pore-size reduction should not be interpreted as the sole mechanism underlying the LDL-dependent prolongation of lysis.

Changes at the level of the fibrin fibers may provide an additional explanation. Fibrin fibers possess distinctive mechanical properties, including high extensibility and strain-dependent stiffness, and fiber mechanics and strain can influence susceptibility to fibrinolytic degradation [3,4]. Prestrain can alter single-fiber lysis behavior [5], while differences in protofibril packing can further influence fibrinolytic susceptibility [6]. LDL could therefore affect lysis through alterations in fibrin organization or material properties that are not fully captured by network pore-size measurements. The present study did not directly measure single-fiber mechanics, fiber molecular packing, or LDL-dependent changes in these properties; consequently, these possibilities remain hypotheses for future investigation.

A further possibility is that LDL directly affects interactions between fibrin and components of the fibrinolytic system. LDL may associate with fibrin or become retained within the developing clot, potentially modifying the local environment available for plasminogen and tPA binding or plasmin generation. Proteomic analyses of plasma clots have demonstrated retention of multiple nonfibrin proteins, including apolipoproteins, within fibrin clots [10]. In parallel, evidence reviewed by Zheng and colleagues suggests extensive cross-talk between lipoprotein metabolism and fibrinolytic mediators, including tPA, PAI-1, plasminogen, and ApoB-containing lipoproteins [20,21]. These observations provide a biological rationale for investigating whether LDL localization within fibrin networks modifies fibrinolytic-protein binding or activity. However, the current experiments did not directly measure LDL–fibrin binding, LDL incorporation into the clot, plasmin generation, or tPA/plasminogen binding. Such mechanisms should therefore be regarded as testable hypotheses rather than conclusions. Colocalization studies using fluorescently labeled LDL, biochemical binding assays, measurements of plasmin generation, and compositional analysis of washed clots could directly address these possibilities.

From a biomaterials perspective, the present findings identify LDL as an endogenous nanoscale constituent capable of modifying both the architecture and degradation behavior of the fibrin scaffold. This concept is consistent with recent work showing that lipid-associated components, including fatty acid–albumin conjugates, can alter fibrin structure, mechanics, and degradation [17]. More broadly, these findings suggest that the physical and functional properties of fibrin clots are not determined solely by fibrinogen and thrombin concentrations but can also be modified by circulating lipid-associated constituents. Such compositional effects may provide a mechanistic bridge between the biochemical environment associated with dyslipidemia and the persistence of thrombi.

At the molecular scale, LDL is not simply a cholesterol carrier but a heterogeneous nanoscale protein–lipid assembly organized around apoB100. Recent structural characterization of human LDL has revealed an extensive apoB100 architecture associated with the lipid particle [9]. This large amphipathic protein–lipid surface provides a plausible physical basis for interactions with fibrin(ogen) or for perturbation of the local physicochemical environment during fibrin polymerization. The present study does not establish whether the observed effects arise from direct apoB100–fibrin interactions, lipid-mediated interactions, or physical incorporation of intact LDL particles into the developing network. Distinguishing among these possibilities will require direct binding, colocalization, and clot-composition measurements.

## 5. Conclusions

Increasing LDL concentration altered plasma fibrin-network architecture and prolonged internal clot lysis across all tested fibrinogen/thrombin conditions. LDL produced a concentration-dependent reduction in network pore size, described by the power-law relationship D_pore = (6.54 ± 0.11)[*LDL*]^−0.12 ±^ 0^.02^, (R^2^ = 0.90; p = 4 × 10^−5^ for the LDL exponent). Internal clot lysis time likewise increased significantly with LDL concentration within each fibrinogen/thrombin/tPA condition, demonstrating a consistent LDL-dependent reduction in fibrinolytic susceptibility. However, the parallel changes in pore size and clot lysis time do not establish a direct causal relationship between network remodeling and impaired fibrinolysis, particularly in light of our previous finding that internal clot lysis can be partially decoupled from thrombin-mediated changes in fibrin architecture [2]. LDL may therefore influence fibrinolysis through additional effects on fibrin organization and material properties, transport and redistribution of fibrinolytic proteins within the network, or direct interactions with components of the fibrinolytic system. Overall, these findings identify LDL as a modifier of both the structural and functional properties of the fibrin biomaterial and suggest a potential mechanism linking elevated LDL to thrombus persistence beyond its established role in atherosclerosis.

## Acknowledgements

This work was supported by National Institutes of Health grants 2R15HL148842-02. The content is solely the responsibility of the authors and does not necessarily represent the official views of the National Institutes of Health. We are grateful to Glen Marrs for his assistance with confocal imaging, and to Daniel Kim-Shaprio for assistance with the absorbance measurements, and to Saeed Movahedi for his assistance with the statistical analysis.

## Author contributions

AN, ADS, SRB, KB, NEH, BEB, MG designed the research, performed experiments, and analyzed the data; AN, CC, MG wrote the manuscript. All authors read and approved the final manuscript.

